# Spatial Signatures of Biological Soil Crusts and Community Level Self-Organization in Drylands

**DOI:** 10.1101/2023.03.21.533724

**Authors:** Daniel Kozar, Bettina Weber, Yu Zhang, Xiaoli Dong

## Abstract

In dryland landscapes, patches of vascular plants can respond to environmental stress by adjusting their spatial pattern to intercept runoff more effectively, i.e., spatially self-organize, and maintain productivity. However, vegetation patch dynamics in drylands often assumes interspaces of plant patches are composed only of bare soil. Biological soil crusts (BSCs) are complex communities, largely of cyanobacteria, algae, lichens, and bryophytes, living in the soil surface in drylands and often cover more area than vascular plants. BSCs often occur in patches of light cyanobacteria and dark-mixed aggregates and can significantly affect and respond to ecohydrological feedbacks in dryland ecosystems. However, little is known about their spatial patterns and dynamics. In this study, we investigate spatial attributes of BSC patches, their spatial interactions with vascular plants, and factors that drive variation in these attributes. We collected ultra-high-resolution (1-cm) data on spatial patterns of BSCs and vascular plants at 26 sites across three ecoregions of the Southwest of the United States of America. Our analysis shows that light cyanobacterial BSCs vary most in their patch shape complexity along the aridity gradient, while dark-mixed BSCs vary strongly in their abundance. The abundance of dark-mixed BSCs is significantly affected by the soil template, namely soil texture and calcareousness, as well as vascular plants to persist under stress. Furthermore, species associations also change with environmental stress. Light cyanobacteria BSCs, likely a significant source of runoff, may act as a buffer for woody plants against drying, as spatial interactions between these biota become more positive (i.e., spatially aggregated) with greater aridity. While dark-mixed BSCs rely significantly on soil conditions and reduce in abundance as a response to aridity stress, we find evidence that they may have some capacity to spatially adjust under conditions of constant aridity. The interaction of dark-mixed BSCs with light cyanobacteria patches becomes more positive with slope. We conclude that light cyanobacteria BSCs can likely change patch shape in response to water limitation, while dark-mixed BSCs have a reduced capacity to do so – providing further evidence that the abundance of dark-mixed BSCs will decline in the future under drying. BSCs and vascular plants coordinate in space in response to resource availability, suggesting the need to consider self-organization of multiple assemblages to fully understand dryland response to climatic change.

## 1 Introduction

Drylands cover approximately 40% of the Earth’s terrestrial surface, are home to roughly an equal proportion of the human population and play key roles in regulating global hydrologic and nutrient cycling (Maestre et al. 2021). As drylands are defined by water limitation, climatic variation can incite significant ecosystem response in these landscapes, and the transition from a productive state to one of bare soil under increased aridity may be abrupt (D’Odorico et al. 2007, Berdugo et al. 2020). As many drylands are projected to experience increased aridity in the future (Cook et al. 2020, Lian et al. 2021), understanding complex ecological responses in these ecosystems is key to estimate risk of aridification and maximize management effectiveness (Wang et al. 2015).

Dryland vegetation, as any sessile organism, often responds to limitation through a reduction in its abundance (total cover), generating a spatial patchiness pervasive to drylands worldwide (von Hardenberg et al. 2010, Okin et al. 2015, Eldridge et al. 2021). However, through ecohydrological feedbacks, patches of vegetation may also persist at the same cover by changing patch shape, rather than size, in response to aridity (Rietkerk et al. 2004, Scheffer 2009, Deblauwe et al. 2012). An increase in connectivity of bare space between vegetation patches as a response to reduced abundance can in turn increase vegetative cover through increased runoff generation (Mayor et al. 2019). Patches of vegetation with adequate dispersal capabilities may therefore receive the same amount of runoff under increased aridity, allowing the ecosystem to maintain vegetative cover within a range of aridity change. Through such ecohydrological feedbacks and patch dynamics, dryland vegetation often self-organizes at large spatial scales – likely increasing ecosystem resilience (Bastiaansen et al. 2018, Rietkerk et al. 2021). As a result, vegetation patches typically become more complex, or less compact, with aridity (Aguiar and Sala 1999). On hillslopes, self-organizing vegetation patches also tend to orient perpendicular to prevailing slope where runoff from interspaces is most readily available (Deblauwe et al., 2012). Spatial self-organization is an important response by sessile organisms to respond to harsh environments, and in drylands, spatial feedbacks mediated by the redistribution of rainfall as runoff by different types of patches play a critical role.

The dominant paradigm of dryland patch dynamics considers only two types of patches, bare soil, and vascular plants (Rietkerk et al. 2004, 2021, Berdugo et al. 2017, Bastiaansen et al. 2018). However, bare soil between vascular plant patches is rarely uninhabited and is often abundant with biota living in the soil surface (Bowker et al. 2018). Biological soil crusts (hereafter BSCs) are complex, largely autotrophic, communities living in the soil surface of drylands between patches of vegetation (Bowker et al. 2013, 2018, Belnap et al. 2016, Chamizo et al. 2016). These poikilohdyric communities often cover more proportional area than vascular plants and play key ecosystem roles through their effects on hydrologic cycling (Bowker et al. 2013, Chamizo et al. 2016, Eldridge et al. 2020, 2021), nitrogen fixation (Barger et al. 2013, 2016, Weber et al. 2015), and soil stabilization (Williams et al. 2012, Belnap and Büdel 2016). In BSCs, filamentous cyanobacteria colonize and stabilize the soil surface via the excretion of exopolysaccharides (EPS) (Colica et al. 2014, Büdel et al. 2016, Kidron et al. 2020, 2022), which promotes the immigration of species with greater water requirements, like dark N-fixing cyanobacteria, lichens, and bryophytes (Belnap et al. 2008, Büdel et al. 2016, Muñoz-Martín et al. 2019, Cantón et al. 2020). While relative abundance, composition, and structure of BSCs vary, they typically form two groups – lightly pigmented cyanobacteria, hereafter light cyanobacteria, and darker aggregated patches of more diverse taxa, hereafter dark-mixed BSCs (Chamizo et al. 2012, Maier et al. 2018, Eldridge et al. 2020, Havrilla et al. 2020). Likely owing to lower water requirements and relatively high lateral growth rates of light cyanobacteria patches, these BSCs often predominate at high levels of abiotic stress compared to dark-mixed BSCs (Read et al. 2016, Sorochkina et al. 2018, Tamm et al. 2018, Becerra-Absalón et al. 2019). Dark-mixed BSC cover has been observed to decrease, sometimes rapidly, in response to increased abiotic stress (Maestre et al. 2015, Ferrenberg et al. 2015). Therefore, BSC groups may have contrasting capacities to adjust patch shape and persist under water limitation – with dark-mixed BSCs likely having a lower capacity to do so.

Spatial patterns of drylands are likely significantly shaped by ecohydrological interactions between BSCs and vascular plants (Kidron 2007, 2018, Zhang et al. 2016, Havrilla et al. 2019, Eldridge et al. 2020, 2021). Light cyanobacteria BSCs are smooth and likely are a source of runoff for vascular plants and dark-mixed BSCs (Eldridge et al. 2000, Yair 2003, Rodríguez-Caballero et al. 2012, Read et al. 2016, Kidron et al. 2020). While the morphological structure and diversity of dark-mixed BSC patches vary, these well-developed crusts increase surface roughness and typically promote pooling and interception of runoff relative to light cyanobacteria BSCs (Kidron 2007, 2019, Chamizo et al. 2016, Guan and Liu 2019). Vascular plants have varying effects on BSCs as they can locally inhibit cyanobacterial growth by deposition of leaf litter (Zhang et al. 2016, Lan et al. 2021), but also promote dark-mixed patches by providing local shading and high soil moisture (Bowker et al. 2016, Navas Romero et al.

2020). These coupled interactions between patches of dryland functional groups likely feedback to significantly affect the redistribution of scarce resources, i.e., water, in drylands (Amir et al. 2022). As such, to understand the spatial patterns of vegetation in drylands and their changes, it is necessary to consider spatial self-organization of multiple interacting groups, that is, spatial self-organization at the community level – beyond the existing theory of spatial self-organization of a single homogenous group.

BSCs, together with vascular plants, form a spatially distributed community and likely organize in space as a response to abiotic stress and local mediation of that stress, creating a dynamic matrix of biota in dryland landscapes. Despite recent developments in mapping BSCs in space with remote sensing technology (Weber and Hill 2016, Rodríguez-Caballero et al. 2017, Rozenstein and Adamowski 2017, Havrilla et al. 2020), little has been done to understand patch dynamics of BSCs and underlying processes driving variations in their patch attributes. In this study, we mapped BSCs across the U.S. Southwest, quantified their patch attributes along the aridity gradient as well as spatial interactions between patches of different functional groups, and lastly, we assessed effects of climatic and environmental drivers on patch attributes and interactions.

## 2 Methods

### 2.1. Field Data Collection

#### 2.1.1. Sites Description

To characterize BSC patches and investigate likely drivers of variation in their characteristics, we selected 26 sites across three ecoregions of drylands in the southwest United States – the Great Basin (9 sites), the Colorado Plateau (9 sites), and the Mojave Desert (8 sites) (Fig. 1; Table S1). These three dryland ecoregions make for an ideal study system as they provide significant variation in environmental and climatic conditions. The Great Basin is a cold desert with frost-heaving and deep soil profiles (Snyder et al. 2019). The Colorado Plateau is also a cold desert with sedimentary soils (Duniway et al. 2016). The Mojave is a hot rain shadow desert characterized by high aridity and nutrient poor alluvial deposits (Thorne 1986). The 26 sites were chosen to cover significant variation in climate, soil, and topography. Climatic water deficit (CWD), a measure of aridity equal to the depth of evaporative demand that exceeds rainfall, ranges between 912 and 1887 mm across sites. Mean site slope varies between 0° and 3.6°, while hillslope aspect includes north and south facing slopes ranging from 0° to 357°. Soil texture at these sites ranges from silty clay loam to sandy loam. All sites are on the property of the United States Bureau of Land Management (BLM), which allots grazing – allowing analysis of the effects of this disturbance.

**Figure 1.**
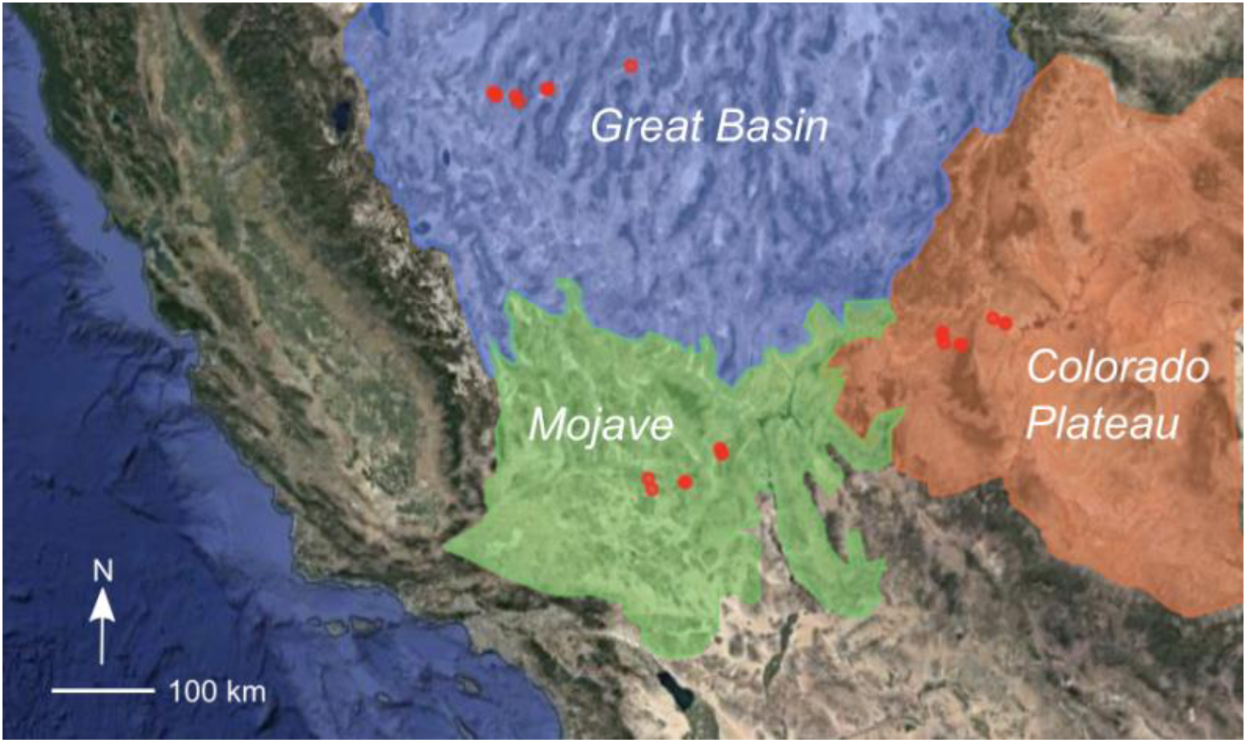
Locations of 26 study sites (red markers) within approximate boundaries of each of the three ecoregions included in this study: the Great Basin, Mojave Desert, and Colorado Plateau. Sites within each region vary by aridity, slope, aspect, and soil conditions (Gorelick et al. 2017).

#### 2.1.2. Community Mapping and Ground Truthing

To map the spatial pattern of BSC-vascular plant communities at each site, we collected 1 cm resolution visible spectrum (RGB) unmanned aerial vehicle (UAV) imagery at all sites in a 50 m by 50 m plot using a DJI Phantom 4 Pro V2.0 at a flight height of 120 feet. UAV flights were conducted from August 16 to August 31 2021 between 10:00 and 15:00 to limit shadow cover. We selected these dates to maximize desiccation of BSCs and limit photosynthetic similarity between them and vascular plants. Collected images were first orthorectified using the photogrammetry software Agisoft Metashape (Agisoft LCC, St. Petersburg, Russia). Orthorectified images were then classified using a maximum likelihood classifier in ENVI version 5.5 (Exelis Visual Information Solutions, Boulder, U.S.A).

We considered a total of eight classes in spectral classification: light cyanobacteria BSC, dark-mixed BSC, woody vegetation, herbaceous vegetation, rock, shadow, non-photosynthetic vegetation, and bare soil. Spectrally unique subclasses of BSCs were considered, namely uniquely colored lichens within dark-mixed BSC patches. All spectral subclasses were combined prior to spatial analysis. 50 ground truthing points were chosen for each class and subclass present to inform the maximum likelihood classifier. 70% of ground truthing points were used for training the classifier, while 30% were used as out-of-sample for cross validation. These ground truthing points were informed from twelve 25 cm by 25 cm quadrats at each site. Large, easily identifiable, woody vegetation patches were not included in quadrats and were identified by images and notes taken at each site. We identified BSC patches and constituent species using a BSC specific field guide (Rosentreter et al. 2007). While some BSCs, typically dark-mixed patches, may grow directly beneath the canopy of woody plants (Zhang et al., 2016), these patches often extend beyond the canopy’s reach at our sites, even in highly arid environments.

The number of spectral subclasses for BSCs was informed by K-means silhouette analysis of three 3m by 3m subplots of BSCs at one site within each ecoregion (Fig. S1). Silhouette analysis identifies the number of spectral classes with the greatest separation distance between them using an unsupervised classification scheme (Shutaywi and Kachouie 2021). Spectral data of BSCs in all regions show greatest divergence when BSCs fall into two classes in our data (Fig. S1), suggesting that these soil communities in our study are constituted of spectrally distinct patches of light cyanobacteria and dark-mixed varieties.

Classification accuracies were assessed with Cohen’s kappa (*k)* coefficients, an overall conservative measure of accuracy across classes (Fitzgerald and Lees 1994), with a minimum *k* of 0.75 to be included in the model (Table S1). We were able to obtain *k* greater than or equal to this cutoff at 22 of the total 26 sites – leaving four sites out of the rest of our analysis. High classification errors at these four sites are likely caused by high photosynthetic similarity between BSCs and vascular plants and movement of ground level objects by high winds.

#### 2.1.3. Soil Sampling

We collected three soil samples of the top 5 cm of soil (Barger et al. 2013, Havrilla and Barger 2018) using a 1 ½” diameter galvanized steel pipe along 25 m intervals at each site. For all soil samples containing BSCs, the top crusted layer was separated from the underlying soil to isolate soil substrate. We then evaluated all soil substrate samples for texture, calcareousness (% as CaCO_3_), and total nitrogen (%) – all considered to significantly affect the composition and abundance of the BSC community (Bryce et al. 2012, Bowker et al. 2016). Soil texture was calculated through the sedimentation method in 100 mL graduated cylinders (Taubner et al. 2009). To validate these measurements, we randomly selected ten soil samples and analyzed soil texture using density analysis (Sheldrick and Wang 1993). We found that these two methods provided statistically similar results (Fig. S2). Soil calcareousness and nitrogen content were obtained through gravimetric loss (U.S. Salinity Laboratory Staff 1954) and combustion methods (AOAC Official Method 972.43 1997), respectively. Soil calcareousness and nitrogen content quantification, as well as the soil texture measurements using density analysis, were all carried out by the UC Davis Analytical Lab.

### 2.2. Data Analysis and Statistical Modelling

#### 2.2.1. Analyses of Spatial Patterns and Principal Components

To characterize the spatial attributes of BSC patches, including both light cyanobacteria and dark-mixed BSCs, we calculated a total of 17 spatial metrics for each BSC group using classified imagery. These 17 metrics quantify patch abundance, patch shape complexity, dominance, edge effects, and aggregation (Table 1). To identify the spatial metrics which capture the most variation in all 17 metrics across sites, we paired principal component analysis (PCA) with linear regression of metric data. PCA reduces data dimensionality by calculating principal component values (PCs) for each site across various metrics, which capture variation in the data with fewer variables (Demšar et al. 2013). High correlation between original metrics and PCs suggests these metrics capture the associated variance in spatial data and allows us to deduce the primary axes of variation for BSC patch traits.

**Table 1.** Spatial metrics for PCA calculated for each BSC patch type.

| Metric | Unit | Description |
| --- | --- | --- |
| Edge density | m ha<sup>-1</sup> | Edge of a class per unit area, measure of complexity |
| Perimeter-area (P-A) ratio | - | Ratio of patch perimeter to area, measure of complexity |
| Coefficient of variation P-A ratio | - | Measure of variance in patch P-A, variation in complexity |
| Aggregation | % | Measure of aggregation of a patch type |
| Proportional cover | - | Measure of total abundance |
| Number of patches | - | Number of unique patches, measure of abundance |
| Mean patch area | ha | Average patch area |
| Clumpiness | - | Measure of aggregation of a patch type |
| Cohesion | % | Connectivity based measure of aggregation of a patch type |
| Division | % | Measure of aggregation of a patch type |
| Mean fractal dimension | - | Average fractal dimension of patches, measure of complexity |
| Interspersion and juxtaposition index (IJI) | % | Measure of intermixing and aggregation of a patch type |
| Largest patch index (LPI) | % | Percent area of largest patch, measure of dominance |
| Landscape shape index (LSI) | - | Measure of aggregation of a patch type |
| Patch density | ha<sup>-1</sup> | Number of patches per unit area, measure of abundance |
| Percent like adjacency (PLADJ) | % | Measure of aggregation of a patch type |
| Edge length | m | Total length of class edge in landscape, measure of complexity |

To ensure that our characterization of spatial attributes is robust to the spatial resolution of aerial imagery, we reduced the resolution of our classified data to 2 cm, half that of the original data and repeated the PCA procedure using the same 17 metrics at each site for each BSC group. To obtain BSC patches at this reduced resolution, new cells were assigned values over a 4 cm^2^ sampling grid (four original cells) in the existing classification raster. If a BSC group occupied less than half of the original cells, the new cell was classified as not occupied by that BSC group. If a group of BSCs occupied half or more of the four original cells within a new sampling grid cell, that new cell was classified as that BSC. If BSC groups occupied equal proportions of the four original cells, light cyanobacteria BSCs were given priority, as this group is understood to be the first developmental stage of BSCs (Read et al. 2016, Sorochkina et al. 2018, Becerra-Absalón et al. 2019). Using these new reduced resolution classifications, we then evaluated PCs of the same 17 metrics. If PCs of BSC metrics at a reduced resolution show similar results as in our 1 cm data, it suggests that the original 1 cm classifications are sufficient to characterize the spatial attributes of patches and do not provide any spurious patchiness.

#### 2.2.2. Patch Orientation

To investigate patch orientation of BSCs, a potential outcome of ecohydrological processes on patch dynamics on hillslopes (Deblauwe et al. 2011, 2012), we evaluated the directional connectivity index (DCI)(Larsen et al. 2012) for each BSC group at each site. DCI determines the connectivity of a patch class, across all patches, at a given angle within a classified matrix (Larsen et al. 2012). We calculated DCI between 0° and 90° from north at 10° intervals to capture all directional variation. With calculated DCI for a given patch class, we then computed the difference between angle of maximum DCI and angle of prevailing slope face of a given site, Δθ (°). As such, Δθ is approximately 90° when patch dominant orientation is perpendicular to the prevailing slope, suggesting the capacity for patches to capture surface runoff by adjusting shape (Deblauwe et al. 2012). However, Δθ likely deviates somewhat from 90° due to microtopography within sites and environmental stochasticity. Therefore, we set 60° as a critical threshold for Δθ to be approximately perpendicular to slope in our analysis. We also expect that Δθ is affected by landscape slope.

#### 2.2.3. Spatial Interactions within Dryland Communities

To characterize the type (positive *vs*. negative) and strength of ecological interaction between dryland functional groups, we calculated the mean spatial interaction, *X*_*i-j*_, between each BSC type and woody vegetation. These interactions include between light cyanobacteria BSCs and woody vegetation, dark-mixed BSCs and woody vegetation, as well as light cyanobacteria and dark-mixed BSCs. While BSCs interact with herbaceous plants as well, this biotic group was significantly present at only 6 of all 22 sites included in analysis. We calculated *X*_*i-j*_, the spatial interaction between groups *i* and *j*, through the spatial analysis through distance indices (SADIE) method developed by Perry 1998. *X*_*i-j*_ is calculated through deriving local spatial interaction from clustering maps (*Eq*. 1)(Perry and Dixon 2002):

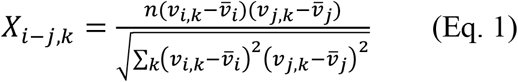

where ν_*i,k*_ *and* ν_*j,k*_ are Perry’s clustering indices (-)(Perry 1998, Perry et al. 1999) of groups *I* and *j* in cell *k* of a 1 m^2^ sampling grid (Navas Romero et al. 2020), and 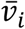 and 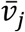 are the mean values of those clustering indices for groups *i* and *j* across all sampling grid cells. Perry’s clustering index *v* of a group in *k*, a unitless metric, is the contribution of *k* to the overall distance to regularity in a sampled abundance grid using proportional cover within each sampling grid cell – the minimum distance that individuals would have to move to obtain an equal abundance across the entire sampling grid (Perry and Dixon 2002). *X*_*i-j,k*_ is the interaction strength between *i* and *j* in *k*, and *n* is the number of cells within the 1 m^2^ sampling grid (*n* = 2500). Local interaction *X*_*i-j, k*_ is increased if both focal groups have ν in *k* greater than or less than the average of the sampling grid, 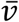; and *X*_*i-j, k*_ is decreased if they deviate from the mean in opposite directions. The numerator of *X*_*i-j, k*_ is normalized through the product of standard deviations of *v* for groups *i* and *j* in the denominator of *Eq*. 1. We calculated a site-specific *X*_*i-j*_ as the mean of *X*_*i-j, k*_ across all sampling grid cells within that site (Perry and Dixon 2002).

*X*_*i-j*_ is particularly useful in this study, as it is standardized to variation in abundance among sites, which allows for comparison of interactions across sites (Perry and Dixon 2002, Navas Romero et al. 2020). *X*_*i-j*_ > 0 indicates positive spatial associations and suggests positive ecological interactions. For example, positive species interactions can result from increased spatial proximity between woody vegetation and dark-mixed BSCs, when vascular plants increase shading for dark-mixed BSCs and dark-mixed BSCs may increase infiltration for those plants (Bowker et al. 2016). *X*_*i-j*_ < 0 indicates negative spatial associations and suggests negative ecological interactions. Competition for water or light between groups, such as burial of light cyanobacteria BSCs by woody vegetation, likely increases spatial segregation of these groups (Zhang et al. 2016, Lan et al. 2021).

#### 2.2.4. Bayesian Statistical Models

To investigate the driving forces of variation in principle spatial attributes of BSC patches, determined from PCA described above, we used a Bayesian multivariate multiple regression. Outcome variables in this model included the first two PCs of each type of BSC patch, light cyanobacteria and dark-mixed BSCs, making four outcome variables in total. Only the first two PCs were used as these two components capture a majority of total variance in spatial metrics, that is, > 65% for both BSC groups across 17 total metrics (*Section 3*). As we expect these four outcome variables might be correlated, we used a multivariate normal distribution for the likelihood function of the model to provide conservative posterior estimates of predictor effect sizes for each outcome:

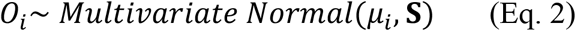

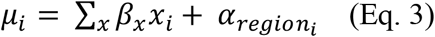

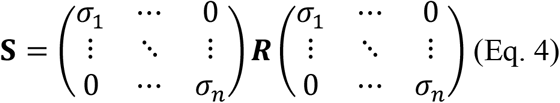

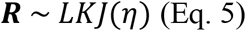

where *n* is the number of outcome variables in the model – *n* = 4 in this case. *β*_*x*_ is the slope parameter describing the effect size of the corresponding predictor variable *x*, including aridity, slope, aspect, grazing, soil texture, soil CaCO_3_, and soil N across all sites (*Eq*. 3). Aridity is represented in our model by climatic water deficit (CWD) (mm) from the TerraClimate dataset, taken as the mean monthly value over the past five years (Abatzoglou et al. 2018). CWD describes water stress through estimation of the total depth of evaporative demand which exceeds available rainfall, with greater values associated with greater aridity. Slope (°), and aspect (°) were extracted from the United States Geological Survey (USGS) 3D Elevation Program (3DEP) dataset (USGS, 2022). Geospatial predictor variables in our model are the average value within the bounds of each site, that is, one value for each variable to represent the mean condition of each site. Predictor data for landscape aspect is converted to northness by taking the cosine of aspect values. Soil texture for individual samples within sites are classified using integers 1-12, increasing in coarseness, according to the United States Department of Agriculture (USDA) system through particle size fractionation analysis. The average of three texture sample values are used to represent soil texture for each site. Soil calcium carbonate (% as CaCO_3_) content and nitrogen content (%) are also averages of the three samples collected within each site. As the data on degree of grazing is scarce and difficult to measure (Fetzel et al. 2017), grazing disturbance is described with a quotient of observed intensity to estimated time since last grazed. This quotient captures the effect of grazing in a continuous variable, as time is expected to reduce the effects of grazing. Grazing intensity at each site was classified 0-3 on a scale of absent – light – moderate – severe, based on our visual inspection in the field (Petz et al. 2014). Light grazing (“1”) is defined as having relatively pristine vegetation with limited evidence of foraging of new vegetative growth. Moderate grazing (“2”) is defined as noticeable foraging of new growth, but still having evidence of original ecosystem structure. Heavy grazing (“3”) shows noticeable evidence of grazing of new growth with evidence of changes to ecosystem structure – such as limited biodiversity and damage to plants (Petz et al. 2014). The time from last major grazing disturbance is classified from current – recent – non-detectible based on evidence of trampling. Time from last major grazing is assigned a “1” if trampling was evidently recent, with disturbed soil showing little recovery. “2” is assigned if trampling was evident but was somewhat recolonized by BSCs. “3” is assigned if disturbed soil is completely recolonized by BSCs, or there is evidence of full ecosystem recovery. This normalization of grazing intensity by a measure of time is necessary as vegetation may not necessarily recover from intense grazing while BSCs may recover in that time. Fire disturbance is not accounted for in our analysis as, according to data from the Monitoring Trends in Burn Severity program, none of our study sites have experienced fire since 1984 (USDA Forest Service and US Geological Survey 2023).

Intercept parameters, *α*, are pooled by region to elucidate local variance in BSC spatial attributes not captured by predictors included in the model. Weak priors are applied to slope parameters *β*_*x*_ and *α*_*region*_. **S** is a variance-covariance matrix for all outcome variables *O*. A Lewandowski-Kurowicka-Joe (LKJ) distribution with shape parameter *η* of 2 was used for a prior distribution for the correlation matrix **R** (*Eqs*. 4 & 5).

In the model described above, we used PC1 and PC2 of the patch spatial metrics of light cyanobacteria BSCs and dark-mixed BSCs as outcome variables. To interpret what each PC represents, we systematically examined the spatial metrics in which these PCs capture the most variance by conducting linear regression analysis between variables. Once we determined the most plausible variable that each PC captures, we built a separate model that is a variant of the one described above. In this model variant, we replaced the PC values on the left side of equation (outcome variables, *O*) (*Eq*. 2) with the values of those most plausible variables we identified. We expect the results of these two models to be largely similar.

We developed a second model to investigate factors driving variation in the interactions between BSC groups and with woody vegetation. The structure of this second model is identical to that of the first model, except the outcome variables are different, reflecting the types of interactions considered. Three types of interactions are included as outcome variables in this model - between light cyanobacteria BSCs and woody vegetation, dark-mixed BSCs and woody vegetation, and light cyanobacteria and dark-mixed BSCs. The two multivariate models share the same set of explanatory variables, including aridity, slope, aspect, grazing, soil texture, soil CaCO_3_ content, and soil N content, as described above.

All data included in our statistical models were standardized by z-scoring to promote model sampling efficiency and allow for direct comparison of effect sizes of different predictors. Posterior distributions of slope and intercept parameters were constructed using 10,000 samples distributed across five chains using Hamiltonian Monte Carlo (HMC) simulation in *Stan* in R (Gelman et al. 2015). These distributions were then used to assess mean standardized slope parameters and degree of confidence in non-zero effect of predictor variables. In this study, central 90% credible intervals are used as a critical degree of confidence to determine statistical significance of parameters. Model convergence was evaluated using trace plots to ensure chains mixed and by ensuring 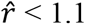 (Gelman and Shalizi 2013).

## 3 Results

Spectral cluster analysis of BSCs in our study suggests that these communities likely fall into two distinct patches – corresponding to light cyanobacteria and dark-mixed BSCs (Fig. S1). BSCs are present at all sites in our study, showing significant variation in proportional cover (Fig. 2A) as well as patch shape (Fig. 2B). Cover of light cyanobacteria BSC ranges from 6% to 58%, while cover of dark-mixed BSC ranges between 0% and 63%. Total coverage of BSCs, including both functional groups, ranges between 11% and 90% across sites. For patches of similar sizes, their shape may vary significantly (Fig. 2B). Mean patch perimeter-area ratio (P-A ratio), a metric describing patch shape complexity, ranges from 1.25 to 1.58 for light cyanobacteria BSCs and 1.30 to 1.54 for dark-mixed BSCs (P-A ratio for individual patches in Fig. 2B, left to right, are 1.66, 2.00, and 2.29). Local interactions with woody vegetation are typically positive for dark-mixed BSCs (mean interaction strength 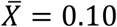 and standard deviation *σ*_*X*_ = 0.28 across sites) (Fig. 2D) and negative for light cyanobacteria BSCs (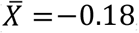, *σ*_*X*_ = 0.21 across sites) (Fig. 2E). The two groups of BSCs are typically well segregated in space (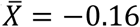, *σ*_*X*_ = 0.26 across sites) (Fig. 2F).

**Figure 2.**
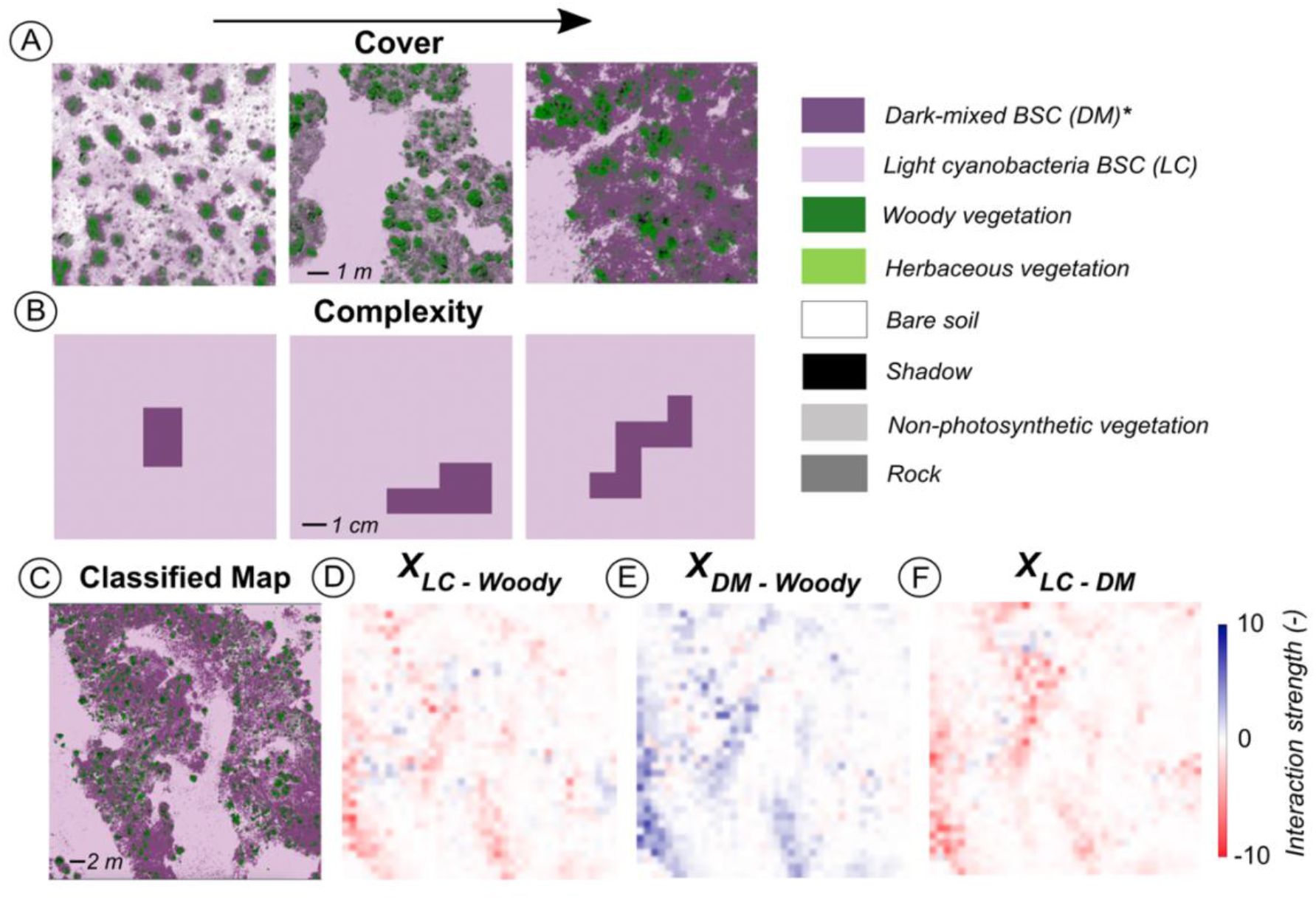
Example classifications displaying variation in (A) abundance and (B) complexity of dark-mixed BSC patches across sites, as well as (C) classified imagery of a site in the Great Basin and (D,E,F) local interactions between woody vegetation and each type of BSC, light cyanobacteria (LC) and dark-mixed (DM) BSCs, within that site.

Through analysis of Δθ, the difference between angle of maximum DCI and angle of dominant hillslope face at a given site, we found that light cyanobacteria BSC patches may tend to orient perpendicular to slope more readily than dark-mixed BSCs (Fig. S3A, B). Although Δθ often deviates from 90° for both BSC groups, the distribution of Δθ for light cyanobacteria tends more toward 90°, with Δθ deviating less than 30° from 90° at 10 sites (Fig. S3A) and Δθ for dark-mixed BSCs is only within 30° deviation from 90° at 5 sites (Fig. S3B). Patch orientation of light cyanobacteria BSCs also exhibits greater dependence on slope, as they orient approximately perpendicular at 6 of 8 sites with slope greater than 2° (Fig. S3A), compared to only 3 sites for dark-mixed BSCs (Fig. S3B).

To further characterize each of the two types of BSC patches, we conducted principal component analysis (PCA) on 17 metrics describing their spatial characteristics (Table 1). The first principal component (PC) of light cyanobacteria BSC patches explained 44.6% of the total variance amongst these 17 metrics, while the second PC explained 29.1% of metric variance (Fig. 3A). We found that the first PC is highly correlated with mean perimeter-area ratio (R^2^ = 0.71, *p* = 0.00) (Fig. 3C), suggesting that this PC captures variation in the complexity of patch shape. The second PC has a significant, high correlation with proportional cover (R^2^ = 0.81, *p* = 0.00) (Fig. 3D), suggesting this PC captures variation in abundance of this BSC type. These results are robust when calculated at a coarser spatial resolution (Fig. S4).

**Figure 3.**
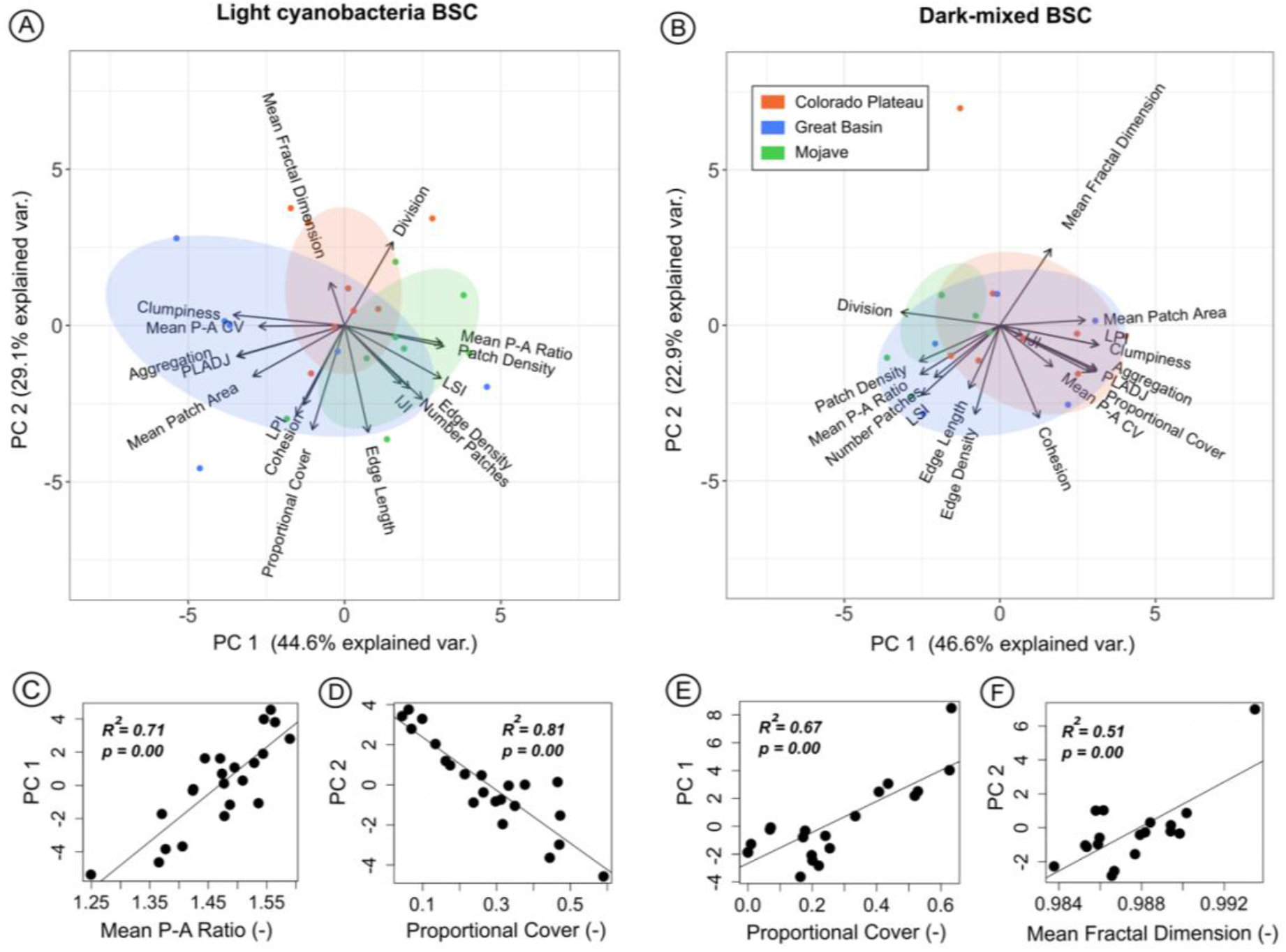
Principal component analysis (PCA) of spatial metrics of BSC patches, showing biplots the first two components for (A) light cyanobacteria BSCs and (B) dark-mixed BSCs. Site specific values of principal components (PCs) are grouped in ellipses by eco-region. Vectors are shown displaying correlation between spatial metrics and both PCs. The x-component of metric vectors corresponds to correlation with PC1, while correlation of spatial metrics with PC2 comprises the y-component of these vectors. (C,D,E,F) Linear relationships between PCs and spatial metrics interpreted to correspond to them.

For dark-mixed BSC patches, PCA analysis showed that the first two principal components of the 17 spatial metrics considered capture 46.6% and 22.9% of the total variance, respectively (Fig. 3B). Unlike light cyanobacteria, the first PC of dark-mixed BSC patches is highly correlated with proportional cover (R^2^ = 0.67, *p* = 0.00) (Fig. 3E) and the second PC is significantly correlated with mean fractal dimension (R^2^ = 0.51, *p* = 0.00) (Fig. 3F) – a measure of patch complexity. Additionally, we found that dark-mixed BSCs vary more significantly in overall abundance than light cyanobacteria, as the standard deviation of proportional cover for these two groups are 0.21 and 0.15, respectively. Like light cyanobacteria BSCs, these results for dark-mixed BSC patch metrics are not significantly changed at a coarser spatial resolution (Fig. S4). The first PC of dark-mixed BSCs at coarser resolution is highly correlated to proportional cover (R^2^ = 0.80, *p* = 0.00) (Fig. S4E) while the second PC at 2 cm is significantly correlated to mean fractal dimension (R^2^ = 0.55, *p* = 0.00) (Fig. S4F). While the correlation between the first two PCs and associated dark-mixed BSC attributes is negative at 2 cm resolution, as opposed to positive at 1 cm, these PCs still likely correspond to the same metrics. Magnitudes of site PC values only differ in sign and are highly correlated between each resolution (Fig. S5). As PCs correspond to eigenvectors of the original metric data, an inverse relationship between PCs and metric still holds the same interpretation as PCs capture the same amount of variance in metric data, and the eigenvectors to which PCs are projected are merely flipped (Bro and Smilde 2014).

We assessed the strength and significance of environmental predictors on variation in the first two principal components of both types of BSC patches through a Bayesian multivariate multiple regression. Model results suggest a statistically significant positive effect of aridity and slope on light cyanobacteria BSC patch complexity (Fig. 4A), with mean effect sizes of 0.72 and 0.61, respectively. Grazing also showed a significant positive effect on light cyanobacteria patch complexity, with a mean effect size of 0.44 (Fig. 4A). For light cyanobacteria BSC cover, soil coarseness showed a significant positive effect, with a mean effect size of 0.56 (Fig. 4B).

**Figure 4.**
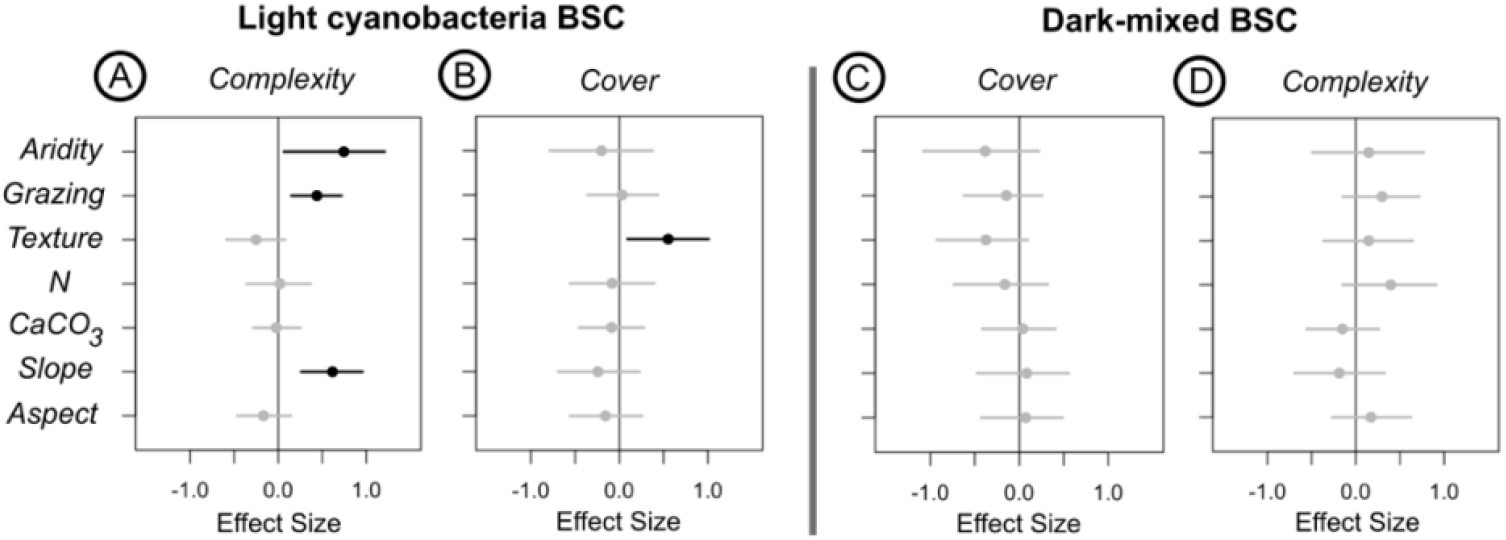
Posterior distributions of predictor effect sizes on (A) light cyanobacteria BSC PC1 (patch complexity) and (B) PC2 (cover) and (C) dark-mixed BSC PC1 (cover) and (D) PC2 (complexity). Mean effect size values are designated by circles and central 90% credible intervals are shown as lines. Parameter distributions statistically different than zero are bolded in black, while those that are insignificantly different from zero are shown in grey.

No predictors included in our analysis exhibited a statistically significant effect on dark-mixed BSC patch complexity or cover (Fig. 4C,D). Aridity and soil texture (coarseness) had the greatest mean effect on the first principal component of dark-mixed BSCs, interpreted as total cover (Fig. 4C). They were marginally significant within 75% and 80% credible intervals, respectively.

These are results of models using PC values as outcome variables. When we directly used the interpreted variables that correspond to each PC as model outcomes (e.g., replacing PC1 with P-A ratio for light cyanobacteria; Fig. 3C), model results stayed largely the same (Fig. S6).

To identify the effect of predictors on variation in interactions between BSC patches and woody vegetation, we specified these interactions as outcome variables in our multivariate regression model. Model results suggest that interaction strengths between BSCs and woody vegetation are significantly affected by aridity, soil texture, and slope. Aridity reduces the degree of spatial segregation (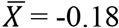 across sites) between light cyanobacteria BSCs and woody vegetation (Fig. 5A). Soil texture (coarseness) enhances the typically positive association between dark-mixed BSCs and woody plants (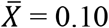 across sites) (Fig. 5B). Soil calcium carbonate (CaCO_3_) however reduces the positive association between dark-mixed BSCs and woody plants (Fig. 5B). The only predictor to show a statistically significant effect on the interaction between light cyanobacteria and dark-mixed BSCs, typically negative (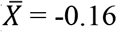 across sites), is slope, which decreases their spatial segregation (Fig. 5C). Aridity also may reduce the spatial segregation between these two groups of BSCs; however, this effect is only marginally significant (Fig. 5C).

**Figure 5.**
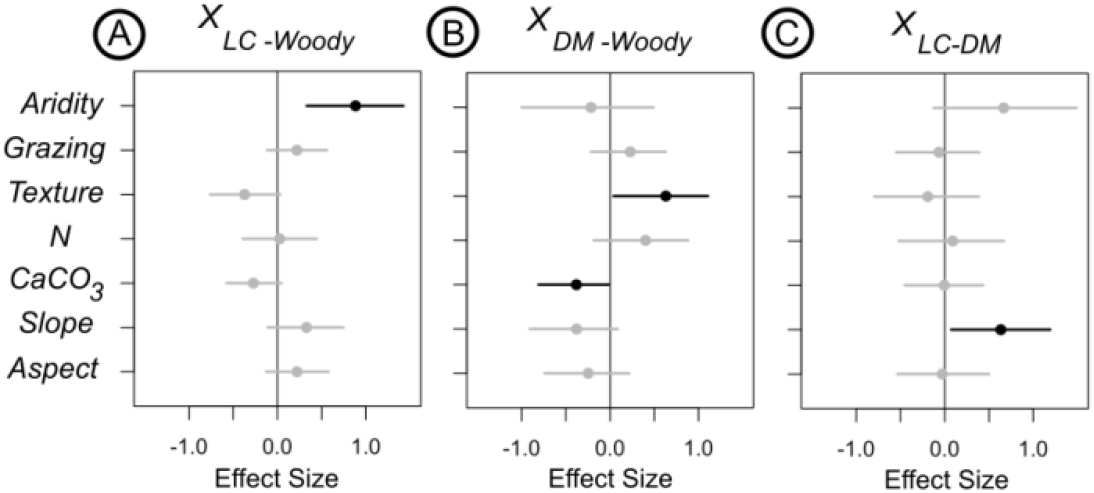
Posterior distributions of predictor effect sizes on the interactions between (A) light cyanobacteria BSC (LC) and woody vegetation, (B) dark-mixed BSC (DM) and woody vegetation, and (C) light cyanobacteria (LC) and dark-mixed BSCs (DM). Mean effect size values are designated by circles and central 90% credible intervals are shown as lines. Parameter distributions statistically different than zero are bolded in black, while those that are insignificantly different from zero are shown in grey.

## 4 Discussion

### 4.1. Traits and Responses of BSC Patches

When the availability of a limiting resource decreases, the total abundance of a sessile organism or functional group will likely decrease in response (Tilman 1982, Svensson and Marshall 2015). However, a patch of biota may receive the same amount of a resource if agents reorganize in space to intercept it more readily (Aguiar and Sala 1999). In drylands, vegetation patches often grow in areas where runoff is easily captured, maintaining productivity under high stress (Aguiar and Sala 1999, Couteron et al. 2014). Organisms with relatively high stress tolerance and relatively rapid growth responses therefore may be able to persist at the same abundance under greater stress through reorganizing in space. We observed such a response in light cyanobacteria BSC patches. The spatial component of greatest variance for light cyanobacteria is patch shape complexity, rather than total cover (Fig. 3A,C), and aridity significantly increases light cyanobacteria patch complexity (Fig. 4A). For dark mixed BSCs, which are more sensitive to aridity (Reed et al. 2012, 2019, Maestre et al. 2015b, Tamm et al. 2018, Ladrón de Guevara and Maestre 2022), the spatial component of greatest variance is cover (Fig. 3B,E), which is likely negatively affected by aridity (Fig. 4C). While the negative effect of aridity is not significant within the 90% credible interval, it is within 75% confidence (Fig. 4C) and a general negative relationship between this BSC group and aridity has also been well documented (Maestre et al. 2015, 2021, Bowker et al. 2016, Read et al. 2016, Weber et al. 2022). Marginal significance of the effect of aridity in our analysis may be caused by differential responses of constituents within dark-mixed BSCs, as lichens and mosses vary both in relative proportion and stress tolerance within well-developed BSC patches (Ferrenberg et al. 2015, Bowker et al. 2016). We also find evidence that light cyanobacterial BSCs patches can more readily orient perpendicular to prevailing hillslope (Fig. S3) — likely also a response of patch reorganization in drylands (Deblauwe et al. 2011). These results suggest that light cyanobacteria-dominated crusts, with greater water stress tolerance and faster lateral growth rate than dark-mixed BSCs (Sorochkina et al. 2018, Becerra-Absalón et al. 2019), can readily adapt to variation in aridity by changing their patch configuration as a response to resource limitation. In contrast, dark-mixed crusts, which are much more sensitive to climatic variation (Reed et al. 2012, Maestre et al. 2015, Rodríguez-Caballero et al. 2022), likely cannot adapt shape sufficiently to stress, resulting in reduced cover under increased aridity.

Sensitive species constituting dark-mixed BSCs have been observed to rely significantly on the physical conditions of soil (Bowker et al. 2016, Weber et al. 2022). We find that coarser substrate likely increases the positive interaction between these BSCs and woody vegetation (Fig. 5B) and plausibly inhibits dark-mixed BSC cover (Fig. 4C). Fine textured soil provides both high water holding capacity and nutrient availability for patches of sensitive dark-mixed BSCs (Bowker et al. 2006, Bowker and Belnap 2008). Under coarser soil conditions, where total cover of dark-mixed BSCs is often reduced (Weber et al. 2022), these organisms likely have greater reliance on local stress amelioration by woody vegetation, which promote shading, relatively high soil moisture, and nutrient availability (Zhang et al. 2016, Havrilla et al. 2019).

While both dark-mixed and light cyanobacteria BSC patches can locally increase soil fineness through deposition (Belnap 2003), our soil texture data capture the mean state of a site and variation across sites. We also find that soil calcareousness likely increases the spatial segregation between woody vegetation and dark-mixed BSCs (Fig. 5B). We speculate that this is likely an effect of lichens. Abundant in calcareous and gypsum soils, where woody vegetation has difficulty establishing (Belnap et al. 2001, Lalley and Viles 2008, Moya et al. 2020), lichens can provide shade and habitat for other sensitive BSC constituents (Rosentreter et al. 2016).

Through increased local shading and soil moisture, lichen abundance may provide functional redundancy at small scales for habitat otherwise provided by woody vegetation (Rosentreter et al. 2016, Havrilla et al. 2019). As lichens have been suggested to themselves have invariant relationships with woody vegetation (Navas Romero et al. 2020), increased abundance of lichens within dark-mixed BSC patches may allow dark-mixed patches persist without woody plants (Fig. 5C). These results highlight that dark-mixed BSC patches likely rely significantly on local soil conditions – essentially binding them in space, and hence, resulting in a reduced capacity to change patch shape to aridity.

### 4.2. Facilitation by Light Cyanobacteria BSCs

Under increasing stress, biota can also persist by shifting in space to reap the benefits of facilitative interactions among species (Kéfi et al. 2016). Light cyanobacterial biocrusts can promote local runoff generation by releasing exopolysaccharides (EPS), which creates a smooth soil surface and clogs soil pores (Chamizo et al. 2016, Kidron et al. 2020, 2022, Eldridge et al. 2020, 2021), likely providing a critical source of runoff for downslope biota (Yair 2001, Rodríguez-Caballero et al. 2018, Eldridge et al. 2020). EPS also likely promotes water adsorption to sediment particles, possibly providing stored water for other biota (Kidron et al. 2020). Our analysis corroborates these findings, showing significant influence of water limitation on the spatial relationships between light cyanobacteria and other dryland biota. Aridity is shown to increase the interaction between woody vegetation and light cyanobacteria BSCs (Fig. 5A). Enhanced runoff through increased slope within a certain range could similarly increase spatial proximity of light cyanobacteria BSCs and woody plants, as slope shows the second greatest mean effect size on this interaction, albeit insignificantly (Fig. 5A). This response is only likely within a range of increased slope before runoff begins to concentrate in rills and vascular plants can no longer effectively intercept runoff by reorganization, possibly explaining the insignificance of this effect in our analysis. These results suggest that under increased water limitation, woody plant patches may shift in space towards light cyanobacteria patches to capture runoff and stored water more effectively. While light cyanobacteria may plausibly shift towards vegetation, this effect is less likely as vegetative cover often inhibits cyanobacteria through deposition of leaf litter (Zhang et al. 2016, Lan et al. 2021). The spatial proximity of dark-mixed BSC patches to light cyanobacteria likely increases as a response to increased slope (Fig. 5C). As we find that dark-mixed BSCs likely lack the capacity to change shape, we expect the spatial interaction between BSC patch types to increase with slope as dark-mixed BSCs most often develop within or adjacent to light cyanobacteria patches. Light cyanobacteria BSCs may develop into dark-mixed BSC patches at favorable microsites where runoff collections, and dark-mixed patches may return to a cyanobacteria-dominant state in unfavorable areas. The mean effect of increased aridity on the interaction between both types of BSCs is less than that of slope and is insignificant, suggesting dark-mixed BSCs have an inhibited capacity to respond to increased climatic stress. Inciting such a patch-level response in runoff sinks of woody vegetation and dark-mixed BSCs suggests the significant role of light cyanobacteria BSCs in organizing spatial structure in drylands and in affecting dryland ecosystem productivity through runoff-runon dynamics (Chamizo et al. 2012a, Rodríguez-Caballero et al. 2018, Eldridge et al. 2020).

### 4.3. Community Level Self-organization in Drylands and Future Work

Biotic agents and their associated spatial feedbacks driven by resource redistribution create a dynamic ecohydrological matrix in drylands (Puigdefábregas 2005, Kéfi et al. 2007, Belnap et al. 2016). Through spatial self-organization and adjustment of spatial patterns, ecosystems can maintain their productivity under increased aridity (Bastiaansen et al. 2018, Rietkerk et al. 2021). In drylands, where abiotic stress is high and likely to increase in many regions, much attention has been given to their responses to decreased water availability, considering only vascular plants (Rietkerk et al. 2004, 2021, Pringle and Tarnita 2017, Bastiaansen et al. 2018). However, our analysis suggests that BSCs, which often cover more area than vascular plants (Bowker et al. 2018), may significantly drive the reorganization of biota in space through the redistribution of scarce water. Patch scale responses of light cyanobacteria BSCs (Fig. 4A), an important source of runoff, to water limitation is paired with spatial adjustments in woody plants (Fig. 5A), and to a limited extent, dark-mixed BSCs (Fig. 5C). In dryland autotrophic communities, high tolerance of light cyanobacteria to stress may therefore act as a buffer for highly sensitive dark-mixed BSCs and vascular plants, which are often considered to be primary runoff sinks (Rodríguez-Caballero et al. 2018). Coupled spatial responses of BSCs and vascular plants to water limitation suggest that the spatial self-organization of dryland ecosystems is likely a community-level response. Changes in ecosystem productivity may be dominated by changes in vascular plant cover; however, BSCs likely significantly alter responses and spatiotemporal dynamics of vascular plants in drylands.

Coordination of spatial responses between multiple functional groups in dryland communities likely significantly affects ecosystem resilience. Responses in tolerant biota, such as light cyanobacteria BSCs, to increasing aridity may offset the incapacity to do so by relatively intolerant BSCs in dark-mixed patches and woody vegetation (Rodríguez-Caballero et al. 2018, Chen et al. 2020). Such a positive interaction might delay an abrupt loss in vegetation cover under increased aridity. While dark-mixed BSCs may alternatively intercept runoff (Rodríguez-Caballero et al. 2012, Chamizo et al. 2016, Guan and Liu 2019), we find they likely lack the capacity to respond to, and persist through, significant increases in stress. Given projected increases in aridity in many drylands (Cook et al. 2020, Lian et al. 2021), several lines of evidence from our study suggest that high successional BSC cover will significantly decrease in coming years, consistent with many other studies (Reed et al. 2012, 2019, Maestre et al. 2015, Ferrenberg et al. 2015, Rodriguez-Caballero et al. 2018). Shading provided by vascular plants and lichens within dark-mixed BSCs may promote persistence of dark-mixed patches, however lichens are also notably susceptible to climate change and disturbance (Navas Romero et al. 2020, Finger-Higgens et al. 2022, Ladrón de Guevara and Maestre 2022). While we find evidence of coupled patch responses of BSCs and vegetation to stress through observational data, controlled experimentation is necessary to fully elucidate mechanisms of patch responses and ecohydrological interactions. Future analysis is also necessary to further characterize spatial responses of dark-mixed BSC patches. Contrasting patch responses of functionally unique biota within dark-mixed BSCs, such as lichens and mosses, may induce inconsistencies in the behavior of dark-mixed patches in aggregate. Our study calls for a new paradigm of considering spatial self-organization in drylands through multiple functional groups — instead of vascular plants only, to improve our predictive capacity of dryland response to change.

## Notes

### Competing Interest Statement

The authors have declared no competing interest.

### Summary of Updates

The manuscript is the same as before, except the authors list has changed so that Dr. Xiaoli Dong is listed last to denote her as my major advisor

## References

Abatzoglou, J. T., S. Z. Dobrowski, S. A. Parks, and K. C. Hegewisch. 2018. TerraClimate, a high-resolution global dataset of monthly climate and climatic water balance from 1958– 2015. Scientific Data 5:170191.

Aguiar, M. R., and O. E. Sala. 1999. Patch structure, dynamics and implications for the functioning of arid ecosystems. Trends in Ecology &Evolution 14:273–277.

Amir, M.-M., S. Ilan, S. Shlomo, A.-G. Hiam, S. Shimshon, and Z. Eli. 2022. The dynamic of the eco-hydrological interrelations between shrubs and biocrusts in the Negev shrublands: Empiric assessments and perspectives for shrubland rehabilitation. CATENA 214:106296.

AOAC Official Method 972.43. 1997. Microchemical Determination of Carbon, Hydrogen, and Nitrogen, Automated Method. Pages 5–6 Official Methods of Analysis of AOAC International. 16th edition. AOAC International, Arlington, VA.

Barger, N. N., S. C. Castle, and G. N. Dean. 2013. Denitrification from nitrogen-fixing biologically crusted soils in a cool desert environment, southeast Utah, USA. Ecological Processes 2:16.

Barger, N.N., Weber, B., Garcia-Pichel, F., Zaady, E., and Belnap, J. 2016. Patterns and Controls on Nitrogen Cycling of Biological Soil Crusts. In: Weber, B., Büdel, B., Belnap, J. (eds) Biological Soil Crusts: An Organizing Principle in Drylands. Ecological Studies, vol 226. Springer, Cham.

Bastiaansen, R., O. Jaïbi, V. Deblauwe, M. B. Eppinga, K. Siteur, E. Siero, S. Mermoz, A. Bouvet, A. Doelman, and M. Rietkerk. 2018. Multistability of model and real dryland ecosystems through spatial self-organization. Proceedings of the National Academy of Sciences 115 (44):11256–11261.

Becerra-Absalón, I., M. Á. Muñoz-Martín, G. Montejano, and P. Mateo. 2019. Differences in the Cyanobacterial Community Composition of Biocrusts From the Drylands of Central Mexico. Are There Endemic Species? Frontiers in Microbiology 10: 937.

Belnap, J. 2003. The World at Your Feet: Desert Biological Soil Crusts. Frontiers in Ecology and the Environment 1(4):181.

Belnap, J., and Büdel, B. 2016. Biological Soil Crusts as Soil Stabilizers. In: Weber, B., Büdel, B., Belnap, J. (eds) Biological Soil Crusts: An Organizing Principle in Drylands. Ecological Studies, vol 226. Springer, Cham

Belnap, J., Büdel, B., and Lange, O.L. 2001. Biological Soil Crusts: Characteristics and Distribution. In: Belnap, J., Lange, O.L. (eds) Biological Soil Crusts: Structure, Function, and Management. Ecological Studies, vol 150. Springer, Berlin, Heidelberg.

Belnap, J., S. L. Phillips, D. L. Witwicki, and M. E. Miller. 2008. Visually assessing the level of development and soil surface stability of cyanobacterially dominated biological soil crusts. Journal of Arid Environments 72:1257–1264.

Belnap, J., Weber, B., and Büdel, B. 2016. Biological Soil Crusts as an Organizing Principle in Drylands. In: Weber, B., Büdel, B., Belnap, J. (eds) Biological Soil Crusts: An Organizing Principle in Drylands. Ecological Studies, vol 226. Springer, Cham

Berdugo, M., M. Delgado-Baquerizo, S. Soliveres, R. Hernández-Clemente, Y. Zhao, J. J. Gaitán, N. Gross, H. Saiz, V. Maire, A. Lehmann, M. C. Rillig, R. v. Solé, and F. T. Maestre. 2020. Global ecosystem thresholds driven by aridity. Science 367:787–790.

Bowker, M. A., and J. Belnap. 2008. A simple classification of soil types as habitats of biological soil crusts on the Colorado Plateau, USA. Journal of Vegetation Science 19:831– 840.

Bowker, M.A. et al. 2016. Controls on Distribution Patterns of Biological Soil Crusts at Micro- to Global Scales. In: Weber, B., Büdel, B., Belnap, J. (eds) Biological Soil Crusts: An Organizing Principle in Drylands. Ecological Studies, vol 226. Springer, Cham.

Bowker, M. A., J. Belnap, D. W. Davidson, and H. Goldstein. 2006. Correlates of biological soil crust abundance across a continuum of spatial scales: support for a hierarchical conceptual model. Journal of Applied Ecology 43:152–163.

Bowker, M. A., D. J. Eldridge, J. Val, and S. Soliveres. 2013. Hydrology in a patterned landscape is co-engineered by soil-disturbing animals and biological crusts. Soil Biology and Biochemistry 61:14–22.

Bowker, M. A., S. C. Reed, F. T. Maestre, and D. J. Eldridge. 2018. Biocrusts: the living skin of the earth. Plant and Soil 429:1–7.

Bryce, S., J. Strittholt, B. Ward, and D. Bachelet. 2012. jColorado Plateau rapid ecoregional assessment report. Denver, USA.

Büdel, B., Dulić, T., Darienko, T., Rybalka, N., and Friedl, T. 2016. Cyanobacteria and Algae of Biological Soil Crusts. In: Weber, B., Büdel, B., Belnap, J. (eds) Biological Soil Crusts: An Organizing Principle in Drylands. Ecological Studies, vol 226. Springer, Cham

Cantón, Y., S. Chamizo, E. Rodriguez-Caballero, R. Lázaro, B. Roncero-Ramos, J. R. Román, and A. Solé-Benet. 2020. Water regulation in cyanobacterial biocrusts from drylands: Negative impacts of anthropogenic disturbance. Water (Switzerland) 12(3).

Chamizo, S., Belnap, J., Eldridge, D.J., Cantón, Y., and Malam Issa, O. (2016). The Role of Biocrusts in Arid Land Hydrology. In: Weber, B., Büdel, B., Belnap, J. (eds) Biological Soil Crusts: An Organizing Principle in Drylands. Ecological Studies, vol 226. Springer, Cham

Chamizo, S., A. Stevens, Y. Cantón, I. Miralles, F. Domingo, and B. van Wesemael. 2012a. Discriminating soil crust type, development stage and degree of disturbance in semiarid environments from their spectral characteristics. European Journal of Soil Science 63:42– 53.

Chamizo, S., Cantón, Y., Rodríguez-Caballero, E., Domingo, F.,& Escudero, A. 2012b. Runoff at contrasting scales in a semiarid ecosystem: A complex balance between biological soil crust features and rainfall characteristics. Journal of Hydrology, 452-453:130–138.

Chen, N., K. Yu, R. Jia, J. Teng, and C. Zhao. 2020. Biocrust as one of multiple stable states in global drylands. Science Advances 6(39).

Colica, G., H. Li, F. Rossi, D. Li, Y. Liu, and R. de Philippis. 2014. Microbial secreted exopolysaccharides affect the hydrological behavior of induced biological soil crusts in desert sandy soils. Soil Biology and Biochemistry 68:62–70.

Cook, B. I., J. S. Mankin, K. Marvel, A. P. Williams, J. E. Smerdon, and K. J. Anchukaitis. 2020. Twenty-First Century Drought Projections in the CMIP6 Forcing Scenarios. Earth’s Future 8.

Couteron, P., F. Anthelme, M. Clerc, D. Escaff, C. Fernandez-Oto, and M. Tlidi. 2014. Plant clonal morphologies and spatial patterns as self-organized responses to resource-limited environments. Philosophical Transactions of the Royal Society A: Mathematical, Physical and Engineering Sciences 372:20140102.

Deblauwe, V., P. Couteron, J. Bogaert, and N. Barbier. 2012. Determinants and dynamics of banded vegetation pattern migration in arid climates. Ecological Monographs 82:3–21.

Deblauwe, V., P. Couteron, O. Lejeune, J. Bogaert, and N. Barbier. 2011. Environmental modulation of self-organized periodic vegetation patterns in Sudan. Ecography 34:990– 1001.

Demšar, U., Harris, P., Brunsdon, C., Fotheringham, A. S., and McLoone, S. 2013. Principal component analysis on Spatial Data: An overview. Annals of the Association of American Geographers, 103(1), 106–128.

Duniway, M. C., T. W. Nauman, J. K. Johanson, S. Green, M. E. Miller, J. C. Williamson, and B. T. Bestelmeyer. 2016. Generalizing Ecological Site Concepts of the Colorado Plateau for Landscape-Level Applications. Rangelands 38:342–349.

Eldridge, D. J., M. Mallen-Cooper, and J. Ding. 2021. Biocrust functional traits reinforce runonrunoff patchiness in drylands. Geoderma 400:115152.

Eldridge, D. J., S. Reed, S. K. Travers, M. A. Bowker, F. T. Maestre, J. Ding, C. Havrilla, E. Rodriguez-Caballero, N. Barger, B. Weber, A. Antoninka, J. Belnap, B. Chaudhary, A. Faist, S. Ferrenberg, E. Huber-Sannwald, O. Malam Issa, and Y. Zhao. 2020. The pervasive and multifaceted influence of biocrusts on water in the world’s drylands. Global Change Biology 26:6003–6014.

Eldridge, D. J., E. Zaady, and M. Shachak. 2000. Infiltration through three contrasting biological soil crusts in patterned landscapes in the Negev, Israel. Catena 40:323–336.

Exelis Visual Information Solutions. (n.d.). ENVI 5.5. Boulder, Colorado.

Ferrenberg, S., S. C. Reed, and J. Belnap. 2015. Climate change and physical disturbance cause similar community shifts in biological soil crusts. Proceedings of the National Academy of Sciences 112:12116–12121.

Fetzel, T., P. Havlik, M. Herrero, and K. Erb. 2017. Seasonality constraints to livestock grazing intensity. Global Change Biology 23:1636–1647.

Finger-Higgens, R., M. C. Duniway, S. Fick, E. L. Geiger, D. L. Hoover, A. A. Pfennigwerth, M. W. van Scoyoc, and J. Belnap. 2022. Decline in biological soil crust N-fixing lichens linked to increasing summertime temperatures. Proceedings of the National Academy of Sciences 119(16): e2120975119.

Fitzgerald, R. W., and B. G. Lees. 1994. Assessing the classification accuracy of multisource remote sensing data. Remote Sensing of Environment 47:362–368.

Gelman, A., D. Lee, and J. Guo. 2015. Stan. Journal of Educational and Behavioral Statistics 40:530–543.

Gelman, A., and C. R. Shalizi. 2013. Philosophy and the practice of Bayesian statistics. British Journal of Mathematical and Statistical Psychology 66:8–38.

Guan, H., and X. Liu. 2019. Does biocrust successional stage determine the degradation of vascular vegetation via alterations in its hydrological roles in semi-arid ecosystem? Ecohydrology 12:1–9.

Gorelick, N., Hancher, M., Dixon, M., Ilyushenko, S., Thau, D., and Moore, R. 2017. Google Earth Engine: Planetary-scale geospatial analysis for everyone. Remote Sensing of Environment 202:18–27.

von Hardenberg, J., A. Y. Kletter, H. Yizhaq, J. Nathan, and E. Meron. 2010. Periodic versus scale-free patterns in dryland vegetation. Proceedings of the Royal Society B: Biological Sciences 277:1771–1776.

Havrilla, C. A., and N. N. Barger. 2018. Biocrusts and their disturbance mediate the recruitment of native and exotic grasses from a hot desert ecosystem. Ecosphere 9(7):e02361.

Havrilla, C. A., V. B. Chaudhary, S. Ferrenberg, A. J. Antoninka, J. Belnap, M. A. Bowker, D. J. Eldridge, A. M. Faist, E. Huber-Sannwald, A. D. Leslie, E. Rodriguez-Caballero, Y. Zhang, and N. N. Barger. 2019. Towards a predictive framework for biocrust mediation of plant performance: A meta-analysis. Journal of Ecology 107:2789–2807.

Havrilla, C. A., M. L. Villarreal, J. L. DiBiase, M. C. Duniway, and N. N. Barger. 2020. Ultra-high-resolution mapping of biocrusts with Unmanned Aerial Systems. Remote Sensing in Ecology and Conservation 6:441–456.

Kéfi, S., M. Holmgren, and M. Scheffer. 2016. When can positive interactions cause alternative stable states in ecosystems? Functional Ecology 30:88–97.

Kéfi, S., M. Rietkerk, M. van Baalen, and M. Loreau. 2007. Local facilitation, bistability and transitions in arid ecosystems. Theoretical Population Biology 71:367–379.

Kidron, G. J. 2007. Millimeter-scale microrelief affecting runoff yield over microbiotic crust in the Negev Desert. CATENA 70:266–273.

Kidron, G. J. 2019. Biocrust research: A critical view on eight common hydrological-related paradigms and dubious theses. Ecohydrology 12(1):e2061.

Kidron, G. J., L. Lichner, T. Fischer, A. Starinsky, and D. Or. 2022. Mechanisms for biocrust-modulated runoff generation – A review. Earth-Science Reviews 231:104100.

Kidron, G. J., Y. Wang, and M. Herzberg. 2020. Exopolysaccharides may increase biocrust rigidity and induce runoff generation. Journal of Hydrology 588:125081.

van de Koppel, J., and M. Rietkerk. 2004. Spatial Interactions and Resilience in Arid Ecosystems. The American Naturalist 163:113–121.

Ladrón de Guevara, M., and F. T. Maestre. 2022. Ecology and responses to climate change of biocrust-forming mosses in drylands. Journal of Experimental Botany 73:4380–4395.

Lalley, J. S., and H. A. Viles. 2008. Recovery of lichen-dominated soil crusts in a hyper-arid desert. Biodiversity and Conservation 17:1–20.

Lan, S., A. D. Thomas, S. Tooth, L. Wu, and D. R. Elliott. 2021. Effects of vegetation on bacterial communities, carbon and nitrogen in dryland soil surfaces: implications for shrub encroachment in the southwest Kalahari. Science of The Total Environment 764:142847.

Larsen, L. G., J. Choi, M. K. Nungesser, and J. W. Harvey. 2012. Directional connectivity in hydrology and ecology. Ecological Applications 22:2204–2220.

Lian, X., S. Piao, A. Chen, C. Huntingford, B. Fu, L. Z. X. Li, J. Huang, J. Sheffield, A. M. Berg, T. F. Keenan, T. R. McVicar, Y. Wada, X. Wang, T. Wang, Y. Yang, and M. L. Roderick. 2021. Multifaceted characteristics of dryland aridity changes in a warming world. Nature Reviews Earth &Environment 2:232–250.

Maestre, F. T., B. M. Benito, M. Berdugo, L. Concostrina-Zubiri, M. Delgado-Baquerizo, D. J. Eldridge, E. Guirado, N. Gross, S. Kéfi, Y. le Bagousse-Pinguet, R. Ochoa-Hueso, and S. Soliveres. 2021. Biogeography of global drylands. New Phytologist 231:540–558.

Maestre, F. T., C. Escolar, R. D. Bardgett, J. A. J. Dungait, B. Gozalo, and V. Ochoa. 2015. Warming reduces the cover and diversity of biocrust-forming mosses and lichens,and increases the physiological stress of soil microbial communities in a semi-arid Pinus halepensis plantation. Frontiers in Microbiology 6.

Maier, S., A. Tamm, D. Wu, J. Caesar, M. Grube, and B. Weber. 2018. Photoautotrophic organisms control microbial abundance, diversity, and physiology in different types of biological soil crusts. The ISME Journal 12:1032–1046.

Mayor, A. G., S. Bautista, F. Rodriguez, and S. Kéfi. 2019. Connectivity-Mediated Ecohydrological Feedbacks and Regime Shifts in Drylands. Ecosystems 22:1497–1511.

Moya, P., A. Molins, S. Chiva, J. Bastida, and E. Barreno. 2020. Symbiotic microalgal diversity within lichenicolous lichens and crustose hosts on Iberian Peninsula gypsum biocrusts. Scientific Reports 10:14060.

Muñoz-Martín, M. Á., I. Becerra-Absalón, E. Perona, L. Fernández-Valbuena, F. Garcia-Pichel, and P. Mateo. 2019. Cyanobacterial biocrust diversity in Mediterranean ecosystems along a latitudinal and climatic gradient. New Phytologist 221:123–141.

Navas Romero, A. L., M. A. Herrera Moratta, E. Martinez Carretero, R. A. Rodriguez, and B. Vento. 2020. Spatial distribution of biological soil crusts along an aridity gradient in the central-west of Argentina. Journal of Arid Environments 176:104099.

Okin, G. S., M. M. las Heras, P. M. Saco, H. L. Throop, E. R. Vivoni, A. J. Parsons, J. Wainwright, and D. P. Peters. 2015. Connectivity in dryland landscapes: shifting concepts of spatial interactions. Frontiers in Ecology and the Environment 13:20–27.

Perry, J. N. 1998. Measures of Spatial Pattern for Counts. Ecology 79:1008.

Perry, J. N., and P. M. Dixon. 2002. A new method to measure spatial association for ecological count data. Écoscience 9:133–141.

Perry, J. N., L. Winder, J. M. Holland, and R. D. Alston. 1999. Red-blue plots for detecting clusters in count data. Ecology Letters 2:106–113.

Petz, K., R. Alkemade, M. Bakkenes, C. J. E. Schulp, M. van der Velde, and R. Leemans. 2014. Mapping and modelling trade-offs and synergies between grazing intensity and ecosystem services in rangelands using global-scale datasets and models. Global Environmental Change 29:223–234.

Pringle, R. M., and C. E. Tarnita. 2017. Spatial Self-Organization of Ecosystems: Integrating Multiple Mechanisms of Regular-Pattern Formation. Annual Review of Entomology 62:359–377.

Puigdefábregas, J. 2005. The role of vegetation patterns in structuring runoff and sediment fluxes in drylands. Earth Surface Processes and Landforms 30:133–147.

Read, C. F., J. Elith, and P. A. Vesk. 2016. Testing a model of biological soil crust succession. Journal of Vegetation Science 27:176–186.

Reed, S. C., K. K. Coe, J. P. Sparks, D. C. Housman, T. J. Zelikova, and J. Belnap. 2012. Changes to dryland rainfall result in rapid moss mortality and altered soil fertility. Nature Climate Change 2:752–755.

Reed, S. C., M. Delgado-Baquerizo, and S. Ferrenberg. 2019. Biocrust science and global change. New Phytologist 223:1047–1051.

Rietkerk, M., R. Bastiaansen, S. Banerjee, J. van de Koppel, M. Baudena, and A. Doelman. 2021. Evasion of tipping in complex systems through spatial pattern formation. Science 374 (6564):eabj0359.

Rietkerk, M., S. C. Dekker, P. C. de Ruiter, and J. van de Koppel. 2004. Self-Organized Patchiness and Catastrophic Shifts in Ecosystems. Science 305:1926–1929.

Rodríguez-Caballero, E., Belnap, J., Büdel, B., Crutzen, P. J., Andreae, M. O., Pöschl, U., and Weber, B. 2018a. Dryland photoautotrophic soil surface communities endangered by Global Change. Nature Geoscience, 11(3):185-189.

Rodríguez-Caballero, E., Y. Cantón, S. Chamizo, A. Afana, and A. Solé-Benet. 2012b. Effects of biological soil crusts on surface roughness and implications for runoff and erosion. Geomorphology 145–146:81–89.

Rodríguez-Caballero, E. S. Chamizo, B. Roncero-Ramos, R. Román, and Y. Cantón. 2018. Runoff from biocrust: A vital resource for vegetation performance on Mediterranean steppes. Ecohydrology 11:e1977.

Rodríguez-Caballero, E., M. Paul, A. Tamm, J. Caesar, B. Büdel, P. Escribano, J. Hill, and B. Weber. 2017. Biomass assessment of microbial surface communities by means of hyperspectral remote sensing data. Science of The Total Environment 586:1287–1297.

Rodríguez-Caballero, E., Reyes, A., Kratz, A., Caesar, J., Guirado, E., Schmiedel, U., and Weber, B. 2022. Effects of climate change and land use intensification on regional biological soil crust cover and composition in Southern Africa. Geoderma, 406:115508

Rodríguez-Caballero, E., J. R. Román, S. Chamizo, B. Roncero Ramos, and Y. Cantón. 2019. Biocrust landscape-scale spatial distribution is strongly controlled by terrain attributes: Topographic thresholds for colonization in a semiarid badland system. Earth Surface Processes and Landforms 44:2771–2779.

Rosentreter, R., M. A. Bowker, and J. Belnap. 2007. A field guide to biological soil crusts of western U.S. drylands: common lichens and bryophytes. U.S. Government Publishing Office, Denver.

Rosentreter, R., D. J. Eldridge, M. Westberg, L. Williams, and M. Grube. 2016. Structure, Composition, and Function of Biocrust Lichen Communities. Pages 121–138.

Rozenstein, O., and J. Adamowski. 2017. A review of progress in identifying and characterizing biocrusts using proximal and remote sensing. International Journal of Applied Earth Observation and Geoinformation 57:245–255.

Scheffer, M. 2009. Critical Transitions in Nature and Society. Princeton University Press.

Sheldrick, B. H., and C. Wang. 1993. Particle-size Distribution. Pages 499–511 Soil Sampling and Methods of Analysis. Canadian Society of Soil Science, Lewis Publishers, Ann Arbor, MI.

Shutaywi, M., and N. N. Kachouie. 2021. Silhouette Analysis for Performance Evaluation in Machine Learning with Applications to Clustering. Entropy 23(6):759.

Snyder, K., J. Huntington, B. Wehan, C. Morton, and T. Stringham. 2019. Comparison of Landsat and Land-Based Phenology Camera Normalized Difference Vegetation Index (NDVI) for Dominant Plant Communities in the Great Basin. Sensors 19(5):1139.

Sorochkina, K. S. Velasco Ayuso, and F. Garcia-Pichel. 2018. Establishing rates of lateral expansion of cyanobacterial biological soil crusts for optimal restoration. Plant and Soil 429:199–211.

Svensson, J. R., and D. J. Marshall. 2015. Limiting resources in sessile systems: food enhances diversity and growth of suspension feeders despite available space. Ecology 96(3):819–827.

Tamm, A., Caesar, J., Kunz, N., Colesie, C., Reichenberger, H., & Weber, B. 2018. Ecophysiological properties of three biological soil crust types and their photoautotrophs from the succulent karoo, South Africa. Plant and Soil, 429(1-2):127–146.

Taubner, H., B. Roth, and R. Tippkötter. 2009. Determination of soil texture: Comparison of the sedimentation method and the laser-diffraction analysis. Journal of Plant Nutrition and Soil Science 172:161–171.

Thorne, R. F. 1986. A Historical Sketch of the Vegetation of the Mojave and Colorado Deserts of the American Southwest. Annals of the Missouri Botanical Garden 73:642.

Tilman, D. 1982. Resource Competition and Community Structure. (MPB-17), Volume 17. Princeton University Press.

Turnbull, L., B. P. Wilcox, J. Belnap, S. Ravi, P. D’Odorico, D. Childers, W. Gwenzi, G. Okin, J. Wainwright, K. K. Caylor, and T. Sankey. 2012. Understanding the role of ecohydrological feedbacks in ecosystem state change in drylands. Ecohydrology 5:174–183.

U.S. Geological Survey. 2022. 3D Elevation Program 10-Meter Resolution Digital Elevation Model.

U.S. Salinity Laboratory Staff. 1954. Alkaline-earth carbonates by gravimetric loss of carbon dioxide. Page 105 Diagnosis and improvement of saline and alkali soils. USDA Agricultural Handbook. U.S. Government Printing Office, Washington, D.C.

USDA Forest Service, and US Geological Survey. 2022. Monitoring Trends in Burn Severity Thematic Burn Severity. Salt Lake City, Utah; Sioux Falls, South Dakota.

Wang, L., S. Manzoni, S. Ravi, D. Riveros-Iregui, and K. Caylor. 2015. Dynamic interactions of ecohydrological and biogeochemical processes in water-limited systems. Ecosphere 6(8):133.

Weber, B., J. Belnap, B. Büdel, A. J. Antoninka, N. N. Barger, V. B. Chaudhary, A. Darrouzet-Nardi, D. J. Eldridge, A. M. Faist, S. Ferrenberg, C. A. Havrilla, E. Huber-Sannwald, O. Malam Issa, F. T. Maestre, S. C. Reed, E. Rodriguez-Caballero, C. Tucker, K. E. Young, Y. Zhang, Y. Zhao, X. Zhou, and M. A. Bowker. 2022. What is a biocrust? A refined, contemporary definition for a broadening research community. Biological Reviews 97:1768–1785.

Weber, B., and Hill, J. 2016. Remote Sensing of Biological Soil Crusts at Different Scales. In: Weber, B., Büdel, B., Belnap, J. (eds) Biological Soil Crusts: An Organizing Principle in Drylands. Ecological Studies, vol 226. Springer, Cham

Weber, B., Wu, D., Tamm, A., Ruckteschler, N., Rodríguez-Caballero, E., Steinkamp, J., Meusel, H., Elbert, W., Behrendt, T., Sörgel, M., Cheng, Y., Crutzen, P.J., Su, H., and Pöschl, U. 2015. Biological soil crusts accelerate the nitrogen cycle through large NO and HONO emissions in drylands. Proc Natl Acad Sci USA. 112(50):15384–9

Williams, A. J., B. J. Buck, and M. A. Beyene. 2012. Biological Soil Crusts in the Mojave Desert, USA: Micromorphology and Pedogenesis. Soil Science Society of America Journal 76:1685–1695.

Yair, A. 2003. Effects of Biological Soil Crusts on Water Redistribution in the Negev Desert, Israel: a Case Study in Longitudinal Dunes. Pages 303–314 in J. Belnap and O. L. Lange, editors. Biological Soil Crusts: Structure, Function, and Management. Springer Berlin Heidelberg, Berlin, Heidelberg.

Zaady, E., I. Katra, H. Yizhaq, S. Kinast, and Y. Ashkenazy. 2014. Inferring the impact of rainfall gradient on biocrusts’ developmental stage and thus on soil physical structures in sand dunes. Aeolian Research 13:81–89.

Zhang, Y., Aradottir, A.L., Serpe, M., and Boeken, B. 2016. Interactions of Biological Soil Crusts with Vascular Plants. In: Weber, B., Büdel, B., Belnap, J. (eds) Biological Soil Crusts: An Organizing Principle in Drylands. Ecological Studies, vol 226. Springer, Cham

