## Supplemental Files for "Spatial Signatures of Biological Soil Crusts and Community Level Self-Organization in Drylands"

***Supplementary Materials***

| Site | Coordinates | Resolution (cm) | Total location error (cm) | Cohen *k* (-) | LC UA (-) | LC PA (-) | DM UA (-) | DM PA (-) |
|---|---|---|---|---|---|---|---|---|
| CP 1 | 112.48°W, 36.94°N | 1.17 | 2.22 | 0.80 | 0.84 | 0.92 | 0.85 | 0.78 |
| CP 2 | 112.49°W, 36.94°N | 1.15 | 1.71 | 0.87 | 0.92 | 0.96 | 0.91 | 0.79 |
| CP 3 | 112.46°W, 36.87°N | 1.17 | 1.09 | 0.93 | 0.93 | 0.84 | 0.82 | 0.91 |
| CP 4 | 111.70°W, 37.07°N | 0.91 | 1.06 | 0.82 | 0.95 | 0.96 | 0.94 | 0.92 |
| CP 5 | 111.70°W, 37.06°N | 0.95 | 1.20 | 0.84 | 0.86 | 0.76 | 0.8 | 0.87 |
| CP 6 | 111.86°W, 37.12°N | 1.14 | 3.04 | 0.88 | 0.93 | 0.95 | 0.93 | 0.74 |
| CP 7 | 112.27°W, 36.86°N | 1.15 | 1.67 | 0.77 | 0.91 | 0.94 | 0.81 | 0.42 |
| CP 8 | 112.26°W, 36.85°N | 1.10 | 1.29 | 0.67 | 0.57 | 0.78 | 0.69 | 0.69 |
| CP 9 | 112.50°W, 36.97°N | 1.12 | 1.30 | 0.94 | 0.95 | 0.95 | 0.96 | 0.91 |
| MJ 1 | 115.25°W, 35.76°N | 1.09 | 1.42 | 0.92 | 0.85 | 0.91 | - | - |
| MJ 2 | 115.25°W, 35.77°N | 1.11 | 1.61 | 0.88 | 0.86 | 0.89 | - | - |
| MJ 3 | 115.28°W, 35.81°N | 1.02 | 1.37 | 0.76 | 0.83 | 0.91 | 0.59 | 0.71 |
| MJ 4 | 115.71°W, 35.47°N | 1.11 | 1.29 | 0.85 | 0.65 | 0.92 | 0.84 | 0.73 |
| MJ 5 | 115.71°W, 35.48°N | 1.07 | 1.36 | 0.89 | 0.89 | 0.94 | 0.92 | 0.92 |
| MJ 6 | 115.71°W, 35.47°N | 1.10 | 1.51 | 0.89 | 0.79 | 0.88 | 0.88 | 0.81 |
| MJ 7 | 116.13°W, 35.40°N | 1.03 | 1.69 | 0.86 | 0.88 | 0.93 | - | - |
| MJ 8 | 116.17°W, 35.50°N | 1.07 | 1.57 | 0.79 | 0.76 | 0.64 | - | - |
| GB 1 | 117.86°W, 39.28°N | 1.06 | 1.46 | 0.97 | 0.98 | 0.98 | 0.91 | 0.98 |
| GB 2 | 117.80°W, 39.24°N | 1.19 | 1.52 | 0.65 | 0.96 | 0.85 | 0.31 | 0.29 |
| GB 3 | 118.09°W, 39.30°N | 1.11 | 1.13 | 0.82 | 0.93 | 0.98 | 0.7 | 0.81 |
| GB 4 | 118.10°W, 39.31°N | 0.95 | 0.94 | 0.85 | 0.86 | 0.94 | 0.61 | 0.86 |
| GB 5 | 118.13°W, 39.33°N | 0.87 | 1.08 | 0.90 | 0.94 | 0.91 | 0.86 | 0.93 |
| GB 6 | 117.46°W, 39.37°N | 1.19 | 1.17 | 0.77 | 0.93 | 0.96 | 0.77 | 0.75 |
| GB 7 | 117.44°W, 39.36°N | 1.05 | 1.12 | 0.90 | 0.95 | 0.98 | 0.82 | 0.89 |
| GB 8 | 117.42°W, 39.36°N | 1.40 | 6.24 | 0.49 | 0.20 | 0.28 | 0.64 | 0.51 |
| GB 9 | 116.41°W, 39.58°N | 1.08 | 4.52 | 0.56 | 0.62 | 0.57 | 0.66 | 0.53 |

**Table S1**. Locations of study sites within the three ecoregions considered in this analysis. Site prefixes denote the ecoregion of each site, with "CP", "MJ", and "GB" corresponding to the Colorado Plateau, Mojave Desert, and Great Basin, respectively. Includes resolution of orthorectified imagery, total horizontal location error from UAV imagery collected, Cohen's kappa coefficient (*k*) for classifications, and the user's accuracy (UA) and producer's accuracy (PA) for light cyanobacteria (LC) and dark-mixed (DM) BSCs. Dark-mixed BSCs were absent from four sites in the Mojave Desert (MJ), where there is no classification accuracy data for this group.

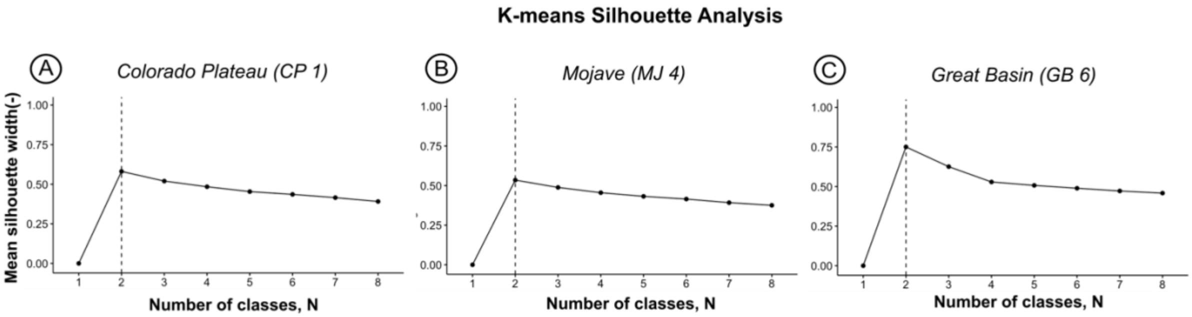


**Figure S1.** Silhouette analysis for BSC spectral classes at sample sites in the three regions in this analysis **-** (a) Colorado Plateau, (b) Mojave Desert and (c) the Great Basin. Greatest mean silhouette width (**-**), a measure of class uniqueness, suggests the optimal number of classes for BSCs, 2 at all regions.

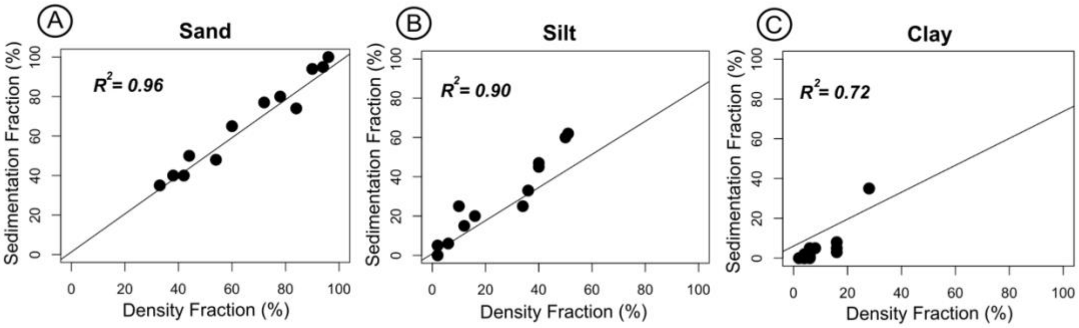


**Figure S2.** Comparison of soil particle fractionation results used to determine texture class between the laboratory hydrometer density samples under solution and the sedimentation method for (a) sand, (b) silt and (c) clay.

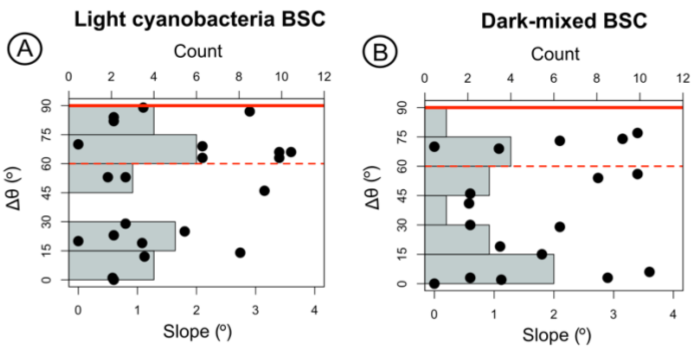


**Figure S3.** The difference between angle of maximum DCI and prevailing site aspect, Δθ (°), plotted against average slope (°) for (A) light cyanobacteria BSC and (B) dark-mixed BSC patches. Included are histograms of Δθ within 15° groupings with total count shown in the x-axes located at the top of each plot. A solid red line is shown at 90°, where Δθ is perpendicular to prevailing aspect, while a dashed red line is shown at 60°, which we set as the lower cutoff for orientation which is approximately perpendicular.

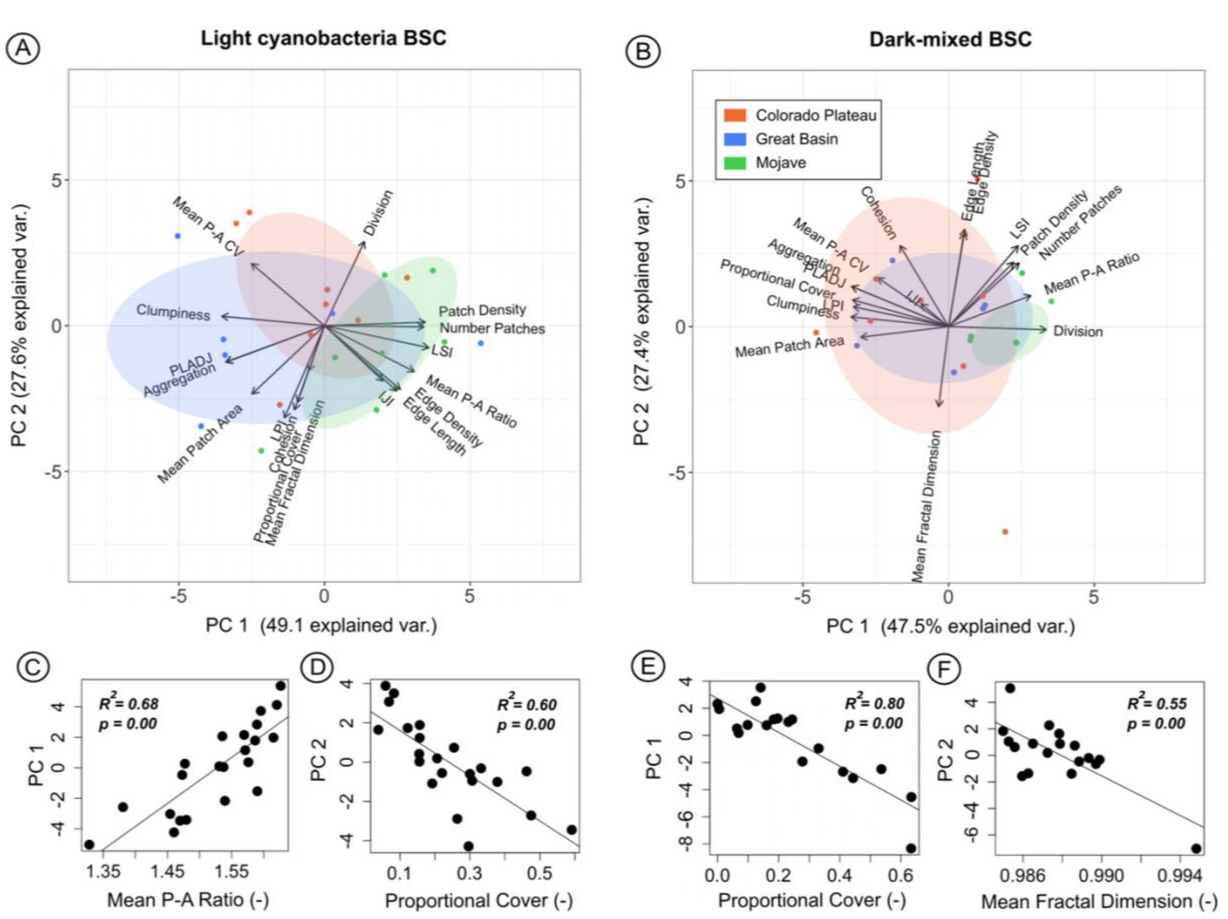


**Figure S4.** Principal component analysis (PCA) showing the first two components for (A) light cyanobacteria BSCs and (B) dark-mixed BSCs at a reduced spatial resolution of 2 cm. Site specific values of principal components (PCs) are grouped by region. Vectors are shown displaying correlation between spatial metrics and both PCs. The x-component of vectors corresponds to correlation with PC1 while correlation of spatial metrics with PC2 comprises the y-component of vectors. (C,D,E,F) Linear regression plots and correlation coefficients between PCs and spatial metrics suspected to correspond to them. The first two PCs of each BSC type show similar correlation to spatial metrics at 2 cm as with the original 1 cm data.

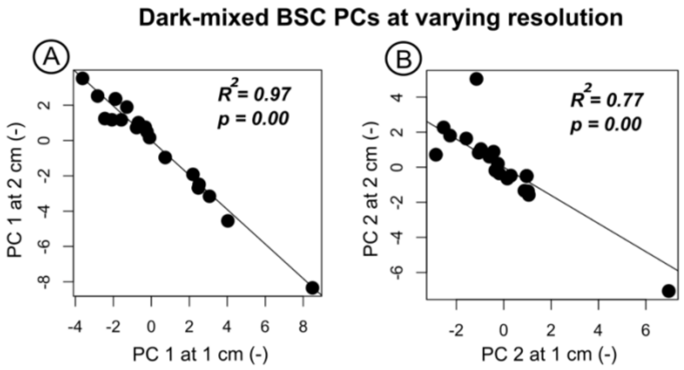


**Figure S5.** Linear correlation plots between (A) the first principal component (PC 1) and (B) the second principal component (PC 2) of dark-mixed BSCs at 1 cm and 2 cm resolution

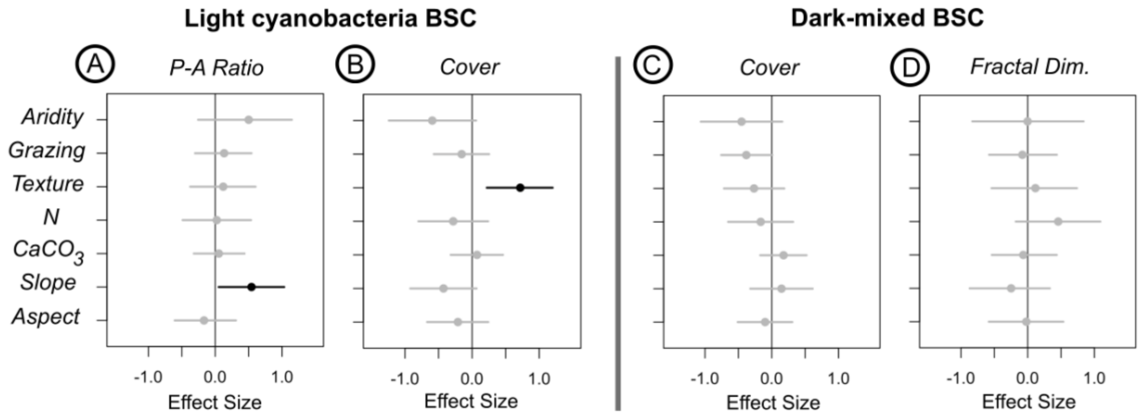


**Figure S6.** Posterior distributions of predictor variable effect sizes on (A) mean light cyanobacteria BSC perimeter to area ratio (P-A ratio) and (B) cover and (C) dark-mixed BSC cover and (D) mean fractal dimension (Fractal Dim.). Mean effect size values are designated by circles. Posterior distributions are within the 90% credible interval, shown as lines. Parameter distributions statistically different than zero are bolded in black, while those that are insignificantly different from zero are shown in light grey.
